# A novel function of *Galactomyces candidum* in highly efficient ammonia nitrogen removal from low C:N wastewater

**DOI:** 10.1101/397687

**Authors:** Li Xiaochao, Liu Hui, Zeng Guihua, Li Hualin, Chen Zhinan, Liu Wenbin, Yang Liping

## Abstract

A strain of bacteria that demonstrated efficient nitrogen removal potential under low C:N conditions was screened from landfill leachate. The strain was identified as *Galactomyces candidum* by ITS sequencing, and growth density and removal of ammonia nitrogen were assessed after 24 h of incubation. The results showed that the optimum ammonia nitrogen reduction conditions for *G. candidum* was at pH 8.0 and 30 °C, with a C:N ratio of 1.5:1; the highest rate of ammonia nitrogen removal was 93.1%. This novel function of *G. candidum* offers great potential in the removal of ammonia nitrogen from sewage, especially in low C:N wastewater. Our study provides a new theoretical basis for the industrial application of bacteria in the biochemical treatment of wastewater and reduces environmental pollution.

A novel function and potential of Galamyces candidum in the removal of ammonia nitrogen from wastewater treatment was found, which provides a theoretical basis for the treatment of ammonia in wastewater.

## Introduction

Removal of ammonia nitrogen from wastewater, especially that with low carbon (C)-nitrogen (N) ratio, is of key interest (You *et al*., 2009; Liang *et al*., 2014). In China, there are high levels of industrial and agricultural pollution in rivers and groundwater deriving from discharge of landfill leachate, petrochemicals, fertilizers, and monosodium glutamate to wastewater that is characterized by low C:N (Jia, 2014). Since traditional biological treatment of wastewater requires large amounts of energy and sources of carbon to complete the denitrification process (Zhao et al.,2001), a more sustainable to approach to wastewater treatment could be the identification of microbes that remain effective under low carbon conditions. Fungi have different carbon-nitrogen ratio requirements, where it is generally >10 (Gao *et al*., 2007). Anammox and denitrifying bacteria have been reported to be two types of autotrophic bacteria effective in low-N:N ratio of sewage (Deng et al., 2015; Persson *et al*., 2017), while Li et al. (2017) and Yang et al. (2018) reported bacterial strains that showed high ammonia nitrogen removal efficiency under low C and N conditions.

*Galactomyces candidum* is a eukaryotic microorganism that is widely present in the environment, has morphological characteristics that are intermediate between yeast and fungus, and is often used in the food and drink industry, including for cheese ripening (Sacristan *et al*., 2012) and production of beer (Sun.2009), white wine, and fermented bean curd, due to the production of pectinase (S.F. Cavalitto, et al., 2006), coenzyme Q, lipase, increase medium chain fatty acid content and antimicrobial activity (Khoramnia, *et al*., 2013). Although *G. candidum* has been reported to reduce the harmful effects of chemical oxygen demand (COD) in sewage treatment of wastewater(He, et al., 2007; Yan, et al.,2002; Xiong, et al.,2008), its impact on the toxic pollutant, ammoniacal nitrogen in landfill leachate is unclear. Here, we screened for strains of *G. candidum* that thrive under low C:N conditions and demonstrated significant effects on ammonia removal from landfill leachate.

## Results and discussion

### Colony morphology

We cultured ammonia nitrogen-reducing strains of *G. candidum* (GC) from landfill leachate in selective medium with NH4+-N as the source of N source. The colony cultured on solid YPD medium at 30 °C was gray-white in color, with a diameter of c. 2 mm, and rough, moist surface, unraised center, and irregular edges (Fig. 1A).

### Strain identification

The GC genome size was approximately 20 kb (Fig. 1B), and ITS sequence lengths of two strains were about 350 and 346 bp (Fig. 1C). BLAST alignment analysis showed similarity between GC and GC strain WM 04.493 KP132255.1 was 99%, and 98% similarity to GC strain LC317625.1. A phylogenetic tree (Kondo *et al*., 2012) of GC strains (Fig. 2) showed differences between GC bacteria and GC strain WM 04.493 KP132255.1 occurred at bases 4, 42, 43, 319, 338, 339, and 340, while differences with GC strain LC317625.1 occurred at bases 65, 91, 92, 319, 338, 339, and 340 (Fig. 3).

**Fig. 1.**
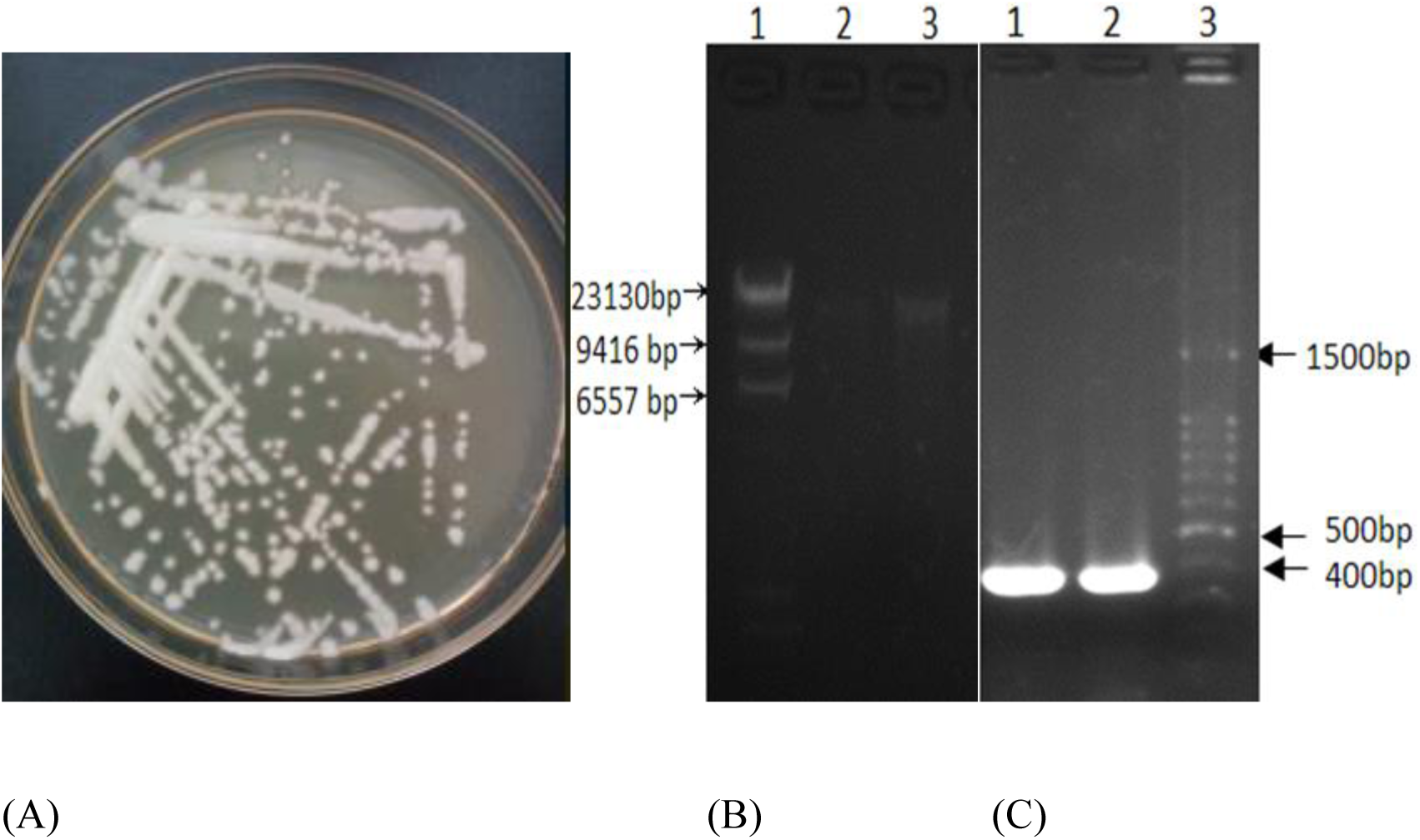
Morphologies of *Galactomyces candidum* colonies cultivated using YPD medium (A); isolation of *G. candidum* strain genome (B), where column 1 shows the DL 23130 DNA markers and columns 2 and 3 show isolation of the *G. candidum* strain genome; identification of the PCR products of *G. candidum* (C), where columns 1 and 2 show PCR products of the ITS rRNA gene and column 3 shows the DL 1500 DNA markers.

**Fig. 2.**
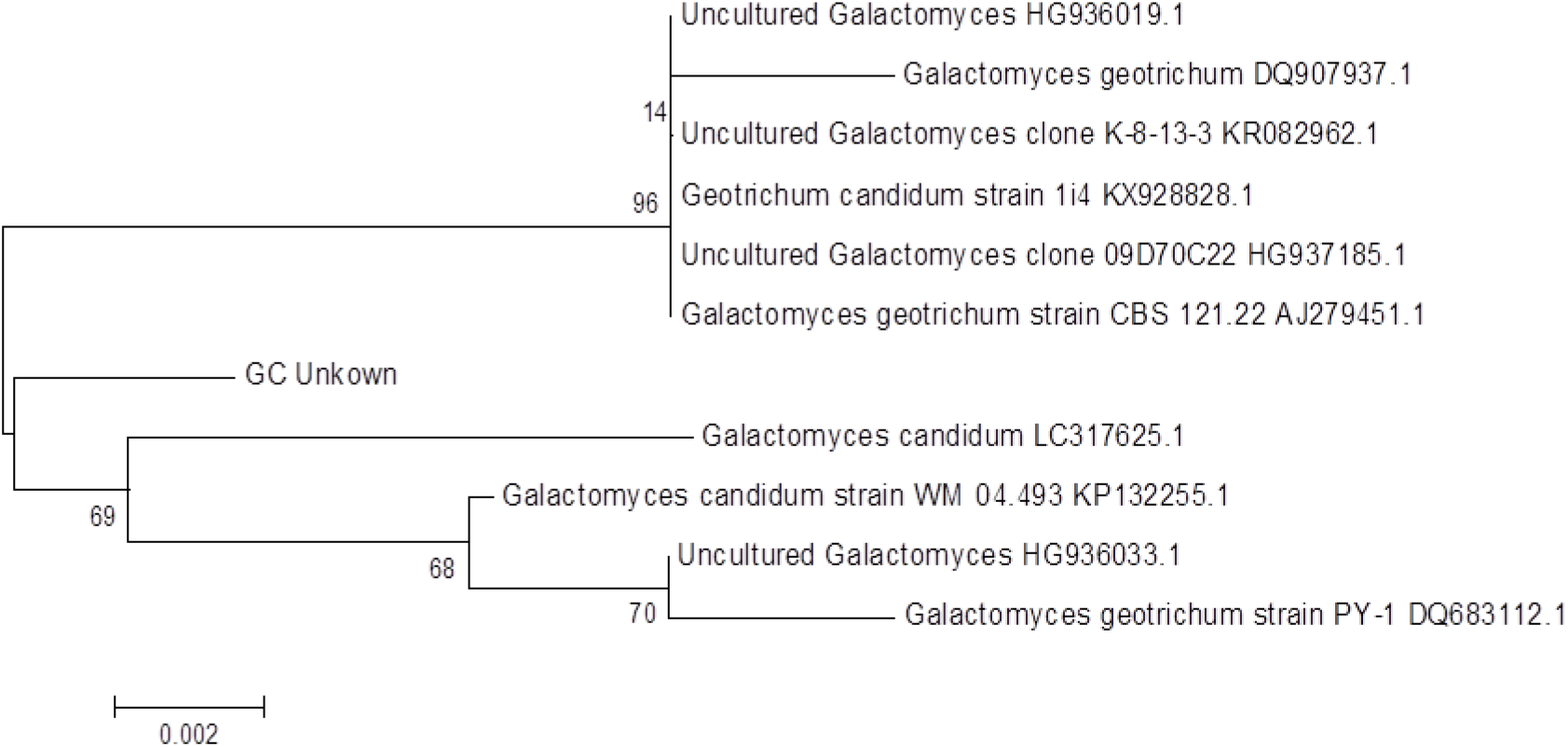
Phylogenetic tree constructed from GC and comparative strains by neighbor-joining method. Bar indicating 0.002 is genetic distance.

**Fig. 3.**
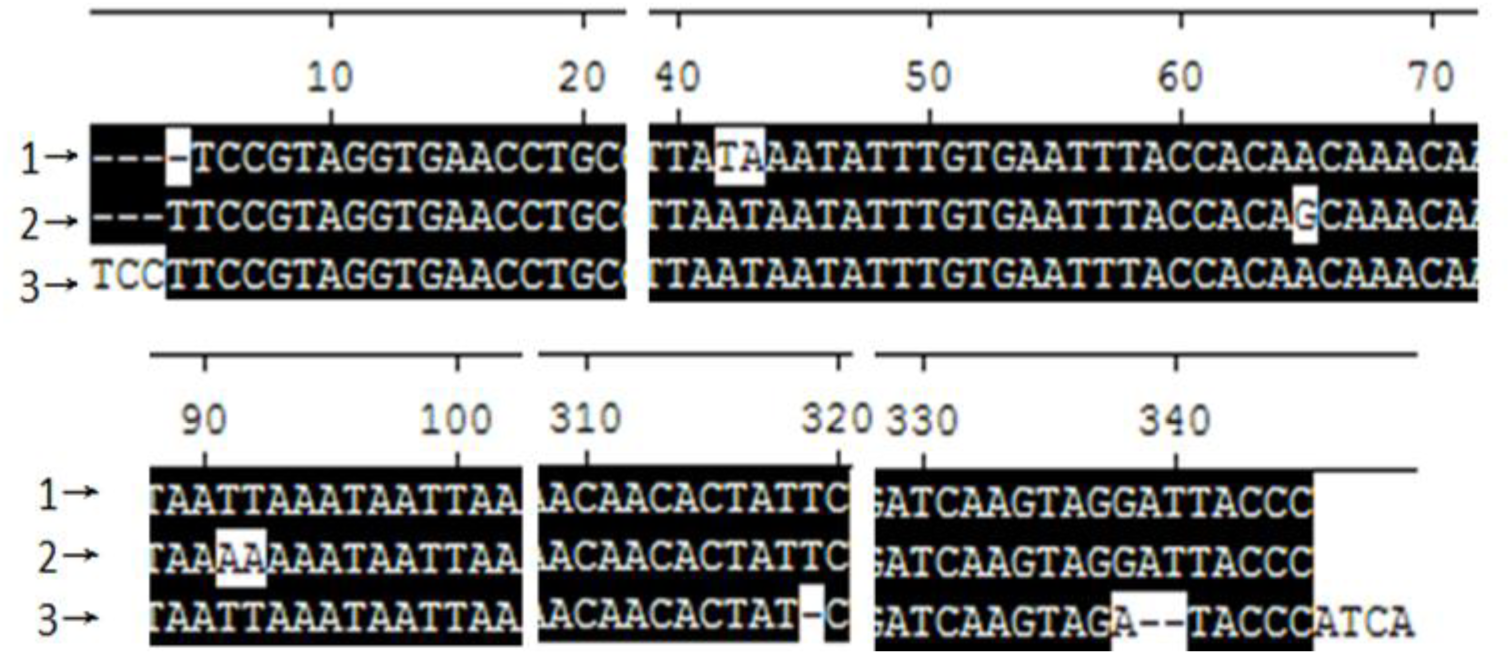
Differences in 16S rRNA gene sequences among *Galactomyces candidum* strains WM 04.493 KP132255.1 (1) and LC317625.1 (2), and GC strains (3).

### GC growth

Growth of *G. candidum* strains was assessed at OD600; the initial delay in growth between 0 and 2 h was followed by logarithmic growth (Fig. 4). This growth pattern showed that the nitrifying bacteria had the characteristics of a short generation cycle with a fast growth rate^[4]^. After 24 h, growth rate was stable. Since turbidity of the bacteria was measured using an ultraviolet spectrophotometer, dead bacteria cells were also counted, and may explain the lack of decline after the stabilization period.

**Fig. 4.**
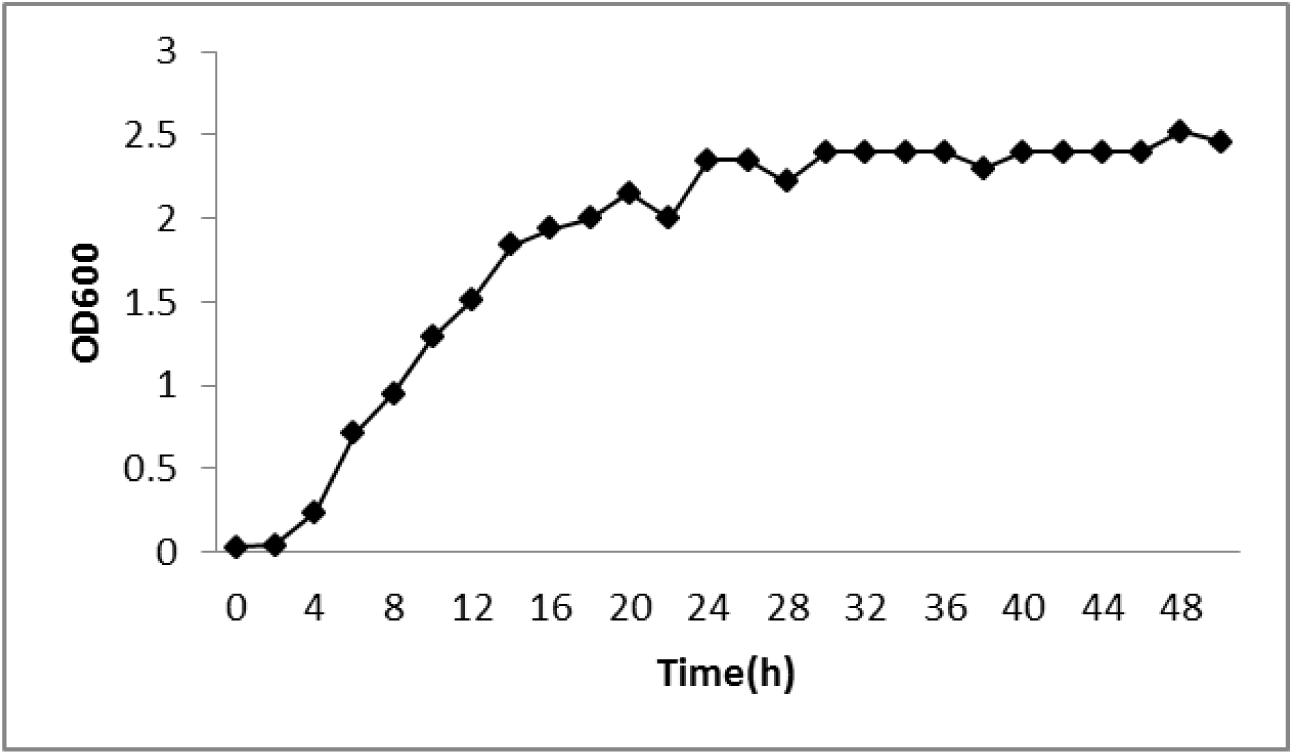
*Galactomyces candidum* growth curve.

### Effect of initial pH on NH _4_ ^+^ -N removal

The optimum pH for growth of *G. candidum* was about 9.0 (Fig. 5A), but ammonia removal efficiency was at pH. 8.0 (Fig. 5B), possibly because the ammonia reducing enzymes in G. candidum are inhibited at high alkaline conditions.

**Fig. 5.**
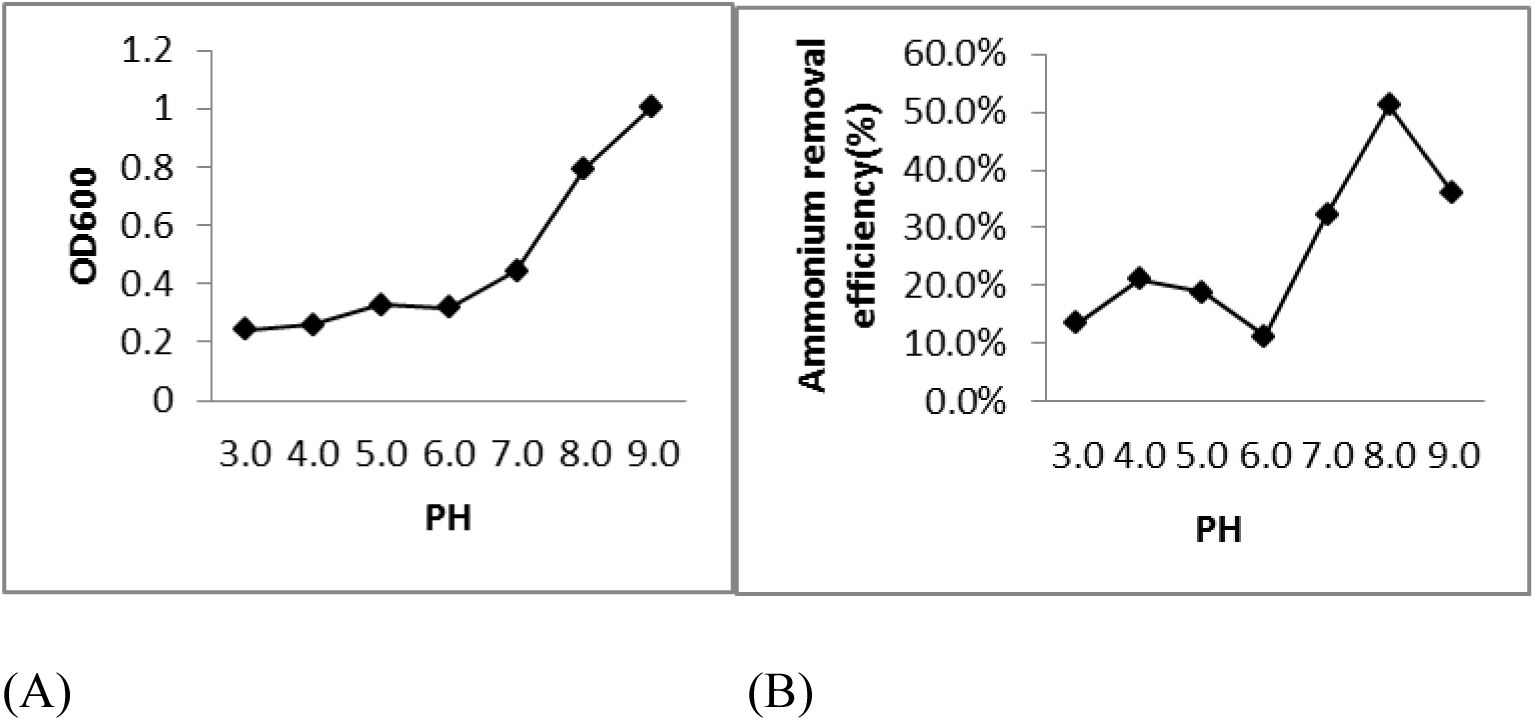
Effect of different pH conditions on *Galactomyces candidum* cell growth (A) and ammonium removal efficiency (B) at 30 °C.

### Effect of temperature on NH _4_ ^+^ -N removal

While *G. candidum* grew at temperatures in the range 10-42 °C, growth was slow at 10 °C and more rapid at temperatures >10 °C (Fig. 6A). We found that the optimum temperature for growth of the bacteria was 30 °C, and growth efficiency decreased when the temperature exceeded 30 °C. Ammonia nitrogen removal rate was highest at 30 °C (76.3%), however this rate reduced to 7% at 42 °C (Fig. 6B). It can be seen that, under different temperature conditions, the optimal growth temperature and ammonia nitrogen removal rate of this strain is 30 °C.

**Fig. 6.**
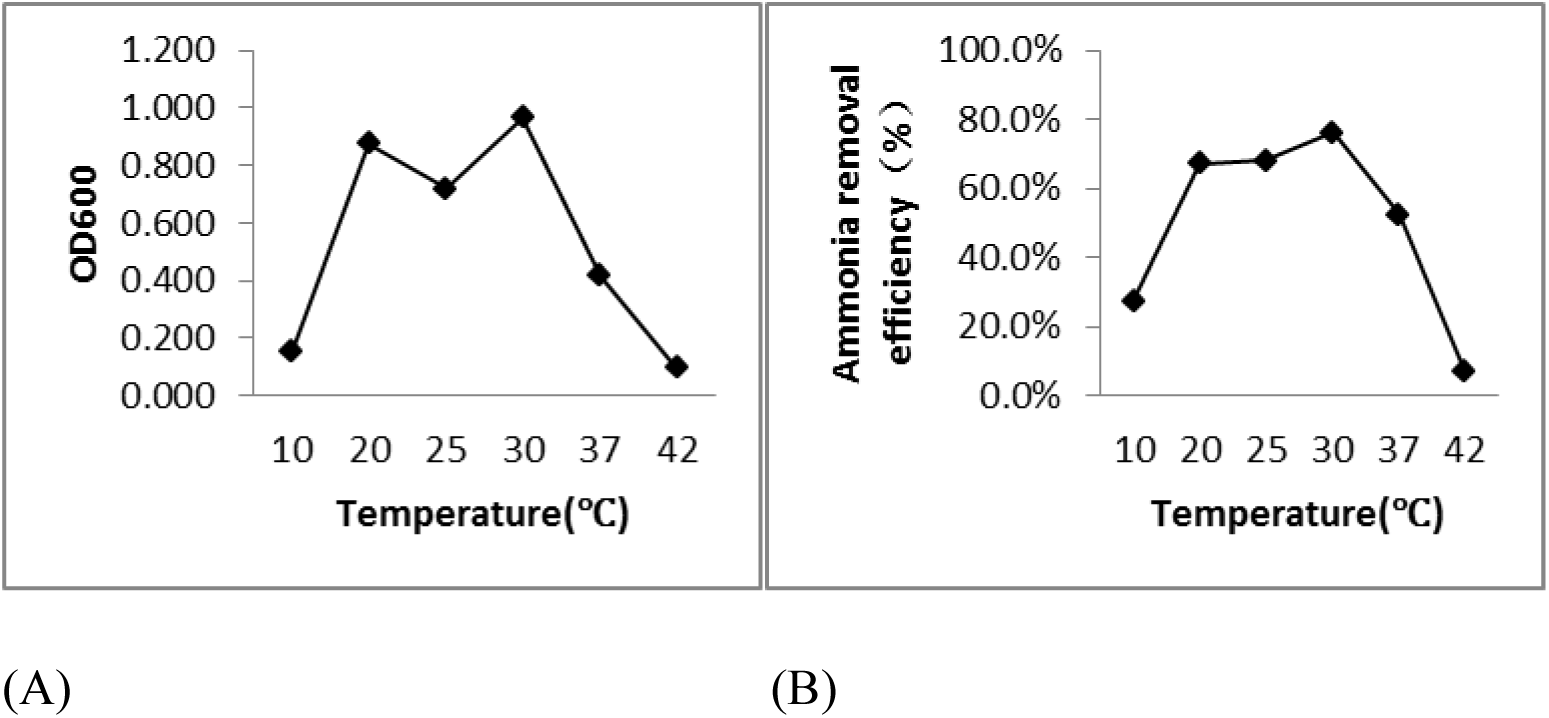
Effect of different temperature conditions on *Galactomyces candidum* cell growth (A) and ammonium removal efficiency (B).

### Effect of C:N ratio on NH_4_ ^+^ -N removal

selection mediumWhen the C:N ratio was low, growth of *G. candidum* was similarly low, but increased as the C increased relative to N (Fig. 7A). Ammonia nitrogen removal rate was similarly low at low C:N ratios (Fig. 7B), possibly as a result of the production of ammonia oxidase by the bacteria that were shown to be a strain of ammoxidation heterotrophs. When the C:N ratio was between 1 and 4, there was an increase in growth of *G. candidum*, with a peak in growth and removal rate of ammonia nitrogen (93.1%) at a C:N ratio of 1.5, and further increases in carbon decreased efficiency of removal of ammonia nitrogen (Fig 7A and 7B).

**Fig. 7.**
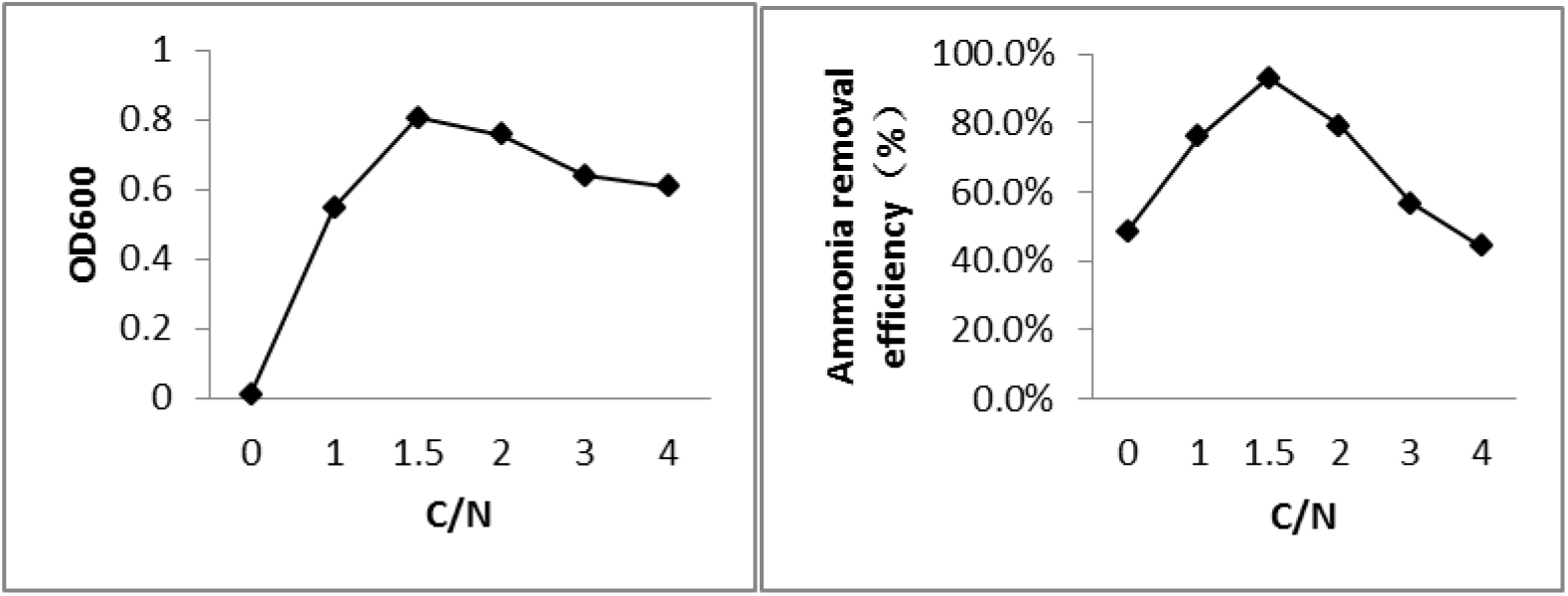
Effect of different C:N ratio conditions on *Galactomyces candidum* cell growth (A) and ammonium removal efficiency (B).

## Conclusions

Industrial wastewater tends to contain high ammonia nitrogen and low C:N ratio, however, insufficient sources of carbons to support traditional biological nitrogen removal results in poor levels of denitrification and the need to increase carbon(Zhu,2015) not only increases economic costs of pollution mitigation, but also increases CO_2_ production and atmospheric pollution. Therefore, a more sustainable approach to wastewater treatment is to maintain the level of carbon use, and explore novel, complementary techniques, but such improvements to the denitrification process may be hampered by various environmental factors (Zhang, 2010). In addition to simultaneous nitrification and denitrification, anaerobic ammonia oxidation and other processes, (Shen, 2013; Zhu, 2016; Peng *et al*., 2018), and new theories and processes have been proposed to reduce the need for carbon in biological denitrification. However, current ammonia nitrogen wastewater treatment that uses strains of microbes to directly reduce the ammonia nitrogen is deemed the most economical and effective. *Galactomyces* screened in this paper grew more rapidly in a neutral to alkaline environment at 20-37 °C, and when the C:N ratio of the medium was 1.5, ammonia nitrogen removal rate was 93.1% after 24 h. Since we found that culturing the bacteria was simple and there was no need to increase carbon, our study provides a theoretical basis for the treatment of ammonia in wastewater.

*Gal. candidum* has effective decolorization and decontamination effects in wastewater treatment (Aouidi *et al*., 2010; Yin.2009) and it has been shown to purify wastewater from the potato, tofu, and corn starch industries(Ren, et al., 2010;Qu, et al., 2005;Li, et al., 2008), however, little is known about the impact of *Gal. candidum* on the removal of ammonia nitrogen from wastewater. We found that the *Galamyces candidum* screened in this paper had a significant effect in the removal of ammonia nitrogen from wastewater after 24 h, especially under high ammonia nitrogen and low carbon conditions, and there have been reports on the use of wastewater to ferment *Galactomyces*. However, it is unclear whether Galactomyces removes ammonia nitrogen via nitrification or denitrification. If used in wastewater treatment, *Galactomyces* could reduce the need for investment in equipment and therefore treatment costs, while efficiently removing ammonia nitrogen.

We found a novel function and potential of *Gal. candidum* in the removal of ammonia nitrogen from wastewater treatment, and it is likely that this bacteria would also elicit positive effects on other pollution factors, such as COD, total nitrogen, and total phosphorus. The effect and safety of use of *Galactomyces* in sewage treatment should be assessed in future research, as should the mechanisms by which ammonia nitrogen is removed by the bacteria.

## Experimental procedures

### Growth conditions

Selection medium (Daum *et al.*, 1998) comprised 0.246 g l^-1^ of NH_4_Cl; 0.039g l^-1^ of MgSO_4_·7H_2_O, 0.055g l^-1^ of CaCl_2_, 0.010g l^-1^ of FeSO_4_·7H_2_O, 2.4 x10^5^ g l^-1^ of CuSO_4_·5H_2_O, 0.348g l^-1^ of K_2_HPO_4_, and 3.603g l^-^1 of glucose. Growth medium (LB) comprised 10.0g l^-1^ of tryptone, 5.0g l^-^1of yeast extract, 10.0g l^-1^ of NaCl, and 20.0g l^-1^ of agar. YPD culture medium comprised 10 g l^-1^ of peptone, 20 g l^-^1of glucose; 5 g l^-1^ of yeast powder, and 20 g l^-1^ of agar powder. MRS isolation medium comprised 10.0 g l^-1^ of peptone, 5.0 g l^-1^ of beef dipping, 4.0 g l^-1^ of yeast dipping powder, 20.0 g l^-1^ of glucose, 2.0 g l^-1^ of K_2_HPO_4_, 2.0 g l^-1^ of C_6_H_17_N_3_O_7_, 5.0 g l^-1^ of CH_3_COONa, 0.2 g l^-1^ of MgSO_4_7H_2_O, 0.05g l^-1^ of MnSO_4_, 20.0g l^-1^ of agar, and 1.0 mL of Tween 80. Media were sterilized at 121 °C for 15 min.

### Strain isolation

We used sterile saline to dilute the landfill leachate that had been collected from a landfill site by 10 times. Glass beads were added to the dilute leachate, which was then placed on a shaker for 20-30 mins to ensure a uniform suspension of microbes. The suspension was strained and transferred to the LB, YPD, and MRS solid media under aseptic conditions using dilution plating, and cultured at 37, 30, and 30 °C, respectively, in an incubator for 24-48 h. Single bacterial strains that formed were selected and further purified by streaking until pure strains were obtained, and then stored at −80 °C. Single colonies of the isolated and purified strains were placed in 5-mL sterile LB, YPD, and MRS liquid media and cultured for 24 h, before 1% of the liquid was removed and centrifuged at 12,000 rpm for 10 min. The liquid was discarded, and then three repeat dilutions using saline were made. Bacteria were added to 100 ml ammoxidation selective medium and placed on a 150 rmp constant shaker in an incubator set at 30 °C for 24 h, and then bacteria were quantified using the turbidity method (OD600). Nessler’s reagent spectrophotometry(HJ 535-2009) was used to measure ammonia nitrogen concentration in the culture medium. Non-inoculated medium was used as a control to calculate the removal rate of ammonia nitrogen, and the strain of *G. candidum* (GC) that had the best effect on ammonia nitrogen removal was selected.

### Bacterial strain identification

Bacteria DNA was extracted from the dilutions using a kit supplied by BIOMIGA (Guangzhou FulenGen Co., Ltd.). We used universal forward ITS1 (5’TCCGTAGGTGAACCTGCGG3’) and reverse ITS4 (5’TCCTCCGCTTATTGATATGC3’) primers to amplify genomic DNA of the isolated strains at 94 °C for 5 min, followed by 30 cycles at 94°C for 0.5 min, 54 °C for 0.75 min, 72 °C for 2 min, 72 °C for 10 min, and then 4 °C. PCR products were sequenced by Tsingke Biological Technology Co., Ltd., Changsha, China. We compared ITS rDNA sequences with those on the NCBI database (https://www.ncbi.nlm.nih.gov/), and a phylogenetic tree was constructed using the neighbor-joining method in MEGA6.0 software.

### Effect of different environmental factors on nitrate reduction

*G. candidum* (GC) bacteria using YPD medium were cultured, and after centrifugation, cells at the bottom of the centrifuge tube were inoculated into ammonia oxidation media. The effect of different initial C:N ratios on ammonia nitrogen removal by GC bacteria cultured at different pH, temperature, and on different media were assessed. Media pH was adjusted to 3.0, 4.0, 5.0, 6.0, 7.0, 8.0, and 9.0, and GC bacteria were cultured in a 150 rmp shaker at 30 °C. Separately, GC bacteria were incubated on a 150 rpm shaker for 24 h at 10, 20, 25, 30, 37, and 42 °C. To assess effect of C:N ratio, we selected ratios equal to 0, 1, 1.5, 2, 3, and 4 and GC bacteria were cultured on 0.246 g of medium fixed with NH4Cl, where glucose was the C source at 30 °C on a 150 rmp shaker. All experiments were conducted using 1% inoculum, with the original medium as a control. Following incubation for 24 h, ammonia nitrogen concentration was assessed at 600 nm OD. All tests comprised three replicates.

### Analytical methods

Cell density was monitored at OD600, using a spectrophotometer, and ammonium nitrogen (NH_4_^+^-N) was measured using Nessler’s reagent spectrophotometry.

## References

[1] You, J., et al. (2009). “Ammonia-oxidizing archaea involved in nitrogen removal.” Water Res 43(7): 1801–1809.

[2] Liang, Y., et al. (2014). “Microbial characteristics and nitrogen removal of simultaneous partial nitrification, anammox and denitrification (SNAD) process treating low C/N ratio sewage.” Bioresour Technol 169: 103–109.

[3] Jia Xiaoning. Study on Low Carbon-to-nitrogen Sewage Denitrification by Combination Bed Based on Immobilized Microorganism [D]. Lanzhou University, 2014.

[4] Zhao Zongsheng, Liu Hongliang, Li Bingwei, Yuan Guangtao. High-efficiency Biological Denitrification of Ammonia Nitrogen Wastewater [J]. CHINA WATER &WASTEWATER,2001(05):24–28.

[5] Gao, L., et al. (2007). “Effects of carbon concentration and carbon to nitrogen ratio on the growth and sporulation of several biocontrol fungi.” Mycol Res 111(Pt 1): 87–92.

[6] Deng Weiwei,Wang Xiaochang. Research progress in the denitrification of sewage at low C/N ratio [J]. Industrial Water Treatment,2015,35(02):15–19

[7] Persson, F., et al. (2017). “Community structure of partial nitritation-anammox biofilms at decreasing substrate concentrations and low temperature.” Microbial Biotechnology 10(4): 761–772.

[8] Li, Y., et al. (2017). “Aerobic-heterotrophic nitrogen removal through nitrate reduction and ammonium assimilation by marine bacterium Vibrio sp. Y1-5.” Bioresour Technol 230: 103–111.

[9] Yang, M., et al. (2018). “Highly efficient nitrogen removal of a coldness-resistant and low nutrient needed bacterium, Janthinobacterium sp. M-11.” Bioresour Technol 256: 366–373.

[10] Sacristan, N., et al. (2012). “Technological characterization of Geotrichum candidum strains isolated from a traditional Spanish goats’ milk cheese.” Food Microbiol 30(1): 260–266.

[11] Sun Bingsheng. Study on Fermentation of Non-alcoholic Beverage Beverage Similar to Beer with Aroma-Producing Strains of GeotrichumCandidum[D]. Shandong Institute of Light Industry,2009.

[12] S.F. Cavalitto,C.F. Mignone. Application of factorial and Doehlert designs for optimization of protopectinase production by a Geotrichum klebahnii strain[J]. Process Biochemistry,2006,42(2).

[13] Khoramnia, A., et al. (2013). “Improvement of medium chain fatty acid content and antimicrobial activity of coconut oil via solid-state fermentation using a Malaysian Geotrichum candidum.” Biomed Res Int 2013: 954542.

[14] He Xiaorong, Hu Shibin,Zheng Zhiwei. Geotrichum candidum for treatment of corn alcohol wastewater and production of single cell protein [J]. Environmental pollution & prevention,2007(10):721-724+730

[15] Yan Bin, Liu Mingdeng. Concentrated Molasses Alcohol Wastewater Fermentation Production of Geotrichum [J]. BIOTECHNOLOGY,2002(03):38–39.

[16] Xiong Zheng, Liu You-xun, Yan Ke-liang, Chen Jing-bo, Zhang Xiao-tong. Study on Fermentation of Geotrichum candidum in High-concentration Inosine Wastewater[J]. Food Science and Technology,2008(03):41–44

[17] Kondo, K., et al. (2012). “Expression, and Molecular and Enzymatic Characterization of Cu-Containing Nitrite Reductase from a Marine Ammonia-Oxidizing Gammaproteobacterium, Nitrosococcus oceani.” Microbes and Environments 27(4): 407–412.

[18] Smith, M.T., Poot, G.A., De Cock, A.W.A.M., 2000. Re-examination of some species of the genus Geotrichum Link:Fr. Antonie van Leeuwenhoek 77,71–8

[19] Oussama Ahrazem, A. P., Juan Antonio Leal (2002). “Fungal cell-wall galactomannans isolated from Geotrichum spp.and their teleomorphs, Dipodascus and Galactomyces.”

[20] Zhu Liang. Research progress on new biological treatment technology for wastewater with high ammonia nitrogen and low C/N ratio[A]. Chinese Society for Environmental Sciences. Proceedings of the 2015 Annual Conference of Chinese Society of Environmental Sciences (Volume Two)[C]. Chinese Society for Environmental Sciences(Chinese Society for Environmental Sciences):,2015:5.

[21] Zhang Shao-hui, Kang Shu-qin. Start-up and Influence Factors of Short cut Nitrification Reactor at Low C/N Ratio [J]. CHINAWATER & WASTEWATER,2010,26(11):96–99

[22] Shen Yanbing. Study on the Sewage Advanced Denitrification Treatment with the low carbon-to-nitrogen ratio [D]. Beijing Jiaotong University,2013

[23] Zhu Tingting. Enhancement of low C/N wastewater treatment in a microbial electrolysis cell[D]. Danlian University of Technology,2016.

[24] Peng, P., et al. (2018). “Effect of adding low-concentration of rhamnolipid on reactor performances and microbial community evolution in MBBRs for low C/N ratio and antibiotic wastewater treatment.” Bioresour Technol 256: 557–561.

[25] Aouidi, F., et al. (2010). “Use of cheese whey to enhance Geotrichum candidum biomass production in olive mill wastewater.” J Ind Microbiol Biotechnol 37(8): 877–882.

[26] Yin Liang. Study of Synergetic Degradation and Decolorization of Azo Dye by Mixed Consortium [D]. South China University of Technology,2009.

[27] Ren Yan, Mei Yun, Zeng Yan, Qin Li-kang. Purification of Wastewater from Potato Starch Production of Ozone Treatment by Geotrichum Candidum [J]. Food Research and Development,2010,31(07):35–38

[28] Qu Jing-ran,Liu Yu,Song Jun-mei. Study on Fermentation of Geotrichum candidum with Waste Water[J]. Food Research and Development,2005(03):99–102.

[29] Li Yu, Li Qi-jiu, Yin Yi-ting, LI Yi-ran,WEI Bo-feng, WANG Xiao-yu, LI Su-yu. The Experimental Condition of Purificationand Being Resources Of Wastewater from Maize Starch Processing by Geotrichum Candidum [J]. Journal of LIAONING University,2008(01):81–84.

[30] Michael Daum, W.Z., 1 Hans Papen,1 Karin Kloos,2 Kerstin Nawrath,2 Hermann Bothe2 “Physiological and Molecular Biological Characterization of Ammonia Oxidation of the Heterotrophic Nitrifier Pseudomonas putida.” Curr.Mierobiol.1998.37(4):281–288.

[31] HJ 535-2009. Water quality. Determination of ammonia nitrogen. Nessler’s reagent spectrophotometry[S].

